# Computational Approaches in Chemical Space Exploration for Carbon Fixation Pathways

**DOI:** 10.1101/2025.09.13.671265

**Authors:** Anne-Susann Abel, Nino Lauber, Jakob Lykke Andersen, Rolf Fagerberg, Daniel Merkle, Christoph Flamm

**Author notes:** **Corresponding Author** Correspondence to Anne-Susann Abel or Christoph Flamm.

## Abstract

Chemical space exploration is an important part of chemistry and biology, enabling the discovery and optimization of metabolic pathways, advancing synthetic metabolic functions, and understanding biochemical network evolution. We use a graph-based computational approach implemented in the cheminformatics software MØD, integrated with Integer Linear Programming (ILP) optimization, to systematically search chemical spaces. This approach allows for flexible and targeted queries, including identification of autocatalytic cycles, thermodynamic considerations, and discovery of novel enzymatic cascades. Specifically, we explore the chemical space of natural and artificial carbon fixation pathways defined from relevant enzyme reactions. By applying different optimization criteria, we identify new varieties and recombinations of natural autocatalytic pathways, and compare the properties of the pathways. This work highlights the versatility of graph-based cheminformatics for the purpose of chemical space exploration and artificial pathway design. Potential applications of this framework extend to carbon capture technologies, improved agricultural yields, and value-added chemical production, advancing efforts to address global sustainability challenges.

## 1 Introduction

As we fight climate change, reducing greenhouse gas emissions to a net zero is crucial. One strategy for this is the removal of CO_2_ from the atmosphere, ideally turning it into value added chemicals for further use [1, 2]. At its core, the process of carbon fixation in living organisms reduces carbon from its most oxidized form (CO_2_ or HCO_3_^2 –^) to valuable metabolic building blocks, e.g. sugars. This is a thermodynamically unfavorable reaction, making large-scale implementation a challenge [1]. However, nature has evolved solutions using enzymatic catalysis and biochemical pathways, coupling more unfavourable reactions to more favourable ones [3, 4]. To date, seven natural carbon fixation pathways have been identified, with several additional artificial pathways proposed [5, 6, 7, 8, 9, 10, 11, 12, 13]. Among the natural pathways, the Acetyl-CoA-Succinyl-CoA pathway family [3] is a particularly interesting group. These include the reductive tricarboxylic acid cycle (rTCA), the dicarboxylate-4-hydroxybutyrate cycle (DC/4-HB), the 3hydroxypropionate-4-hydroxybutyrate cycle (3-HP/4-HB), and the 3-hydroxypropionate bicycle (3-HP bicycle). This group of pathways shares a common structural feature: one half of the cycle converts Succinyl-CoA to Acetyl-CoA, while the other half catalyzes the reverse reaction. Furthermore, each of these pathways overlaps significantly with at least one other, making them highly relevant as templates for designing artificial pathways.

Another commonality between these natural carbon fixation pathways is their autocatalytic nature, which is an important property of several biochemical pathways [14]. A reaction is autocatalytic if at least one reaction product is a catalyst in the reaction producing this product [15]. In an autocatalytic pathway, at least one metabolite, the autocatalyst, acts as a catalyst for its own formation, and together with external inputs, each cycle yields a net gain of this metabolite [14, 16]. This autocatalyst needs to be present for the pathway to start up initially.

Autocatalysis is at the heart of many pathways in the central carbon metabolism [14], and therefore an interesting property to look for when combining reactions of the carbon metabolism from different organisms to design novel pathways, as demonstrated by [17]. Previous advances in artificial pathway design, such as the work of the Erb Lab [18, 19], demonstrate that combining reactions from diverse domains of life and optimizing them for thermodynamic favorability can yield novel, highly efficient pathways. These previous approaches [17, 18, 19] use a combination of heuristic considerations and conceptual analysis, thermodynamic optimization, database searches, and an extensive experimental phase with several steps of optimization, including site-directed mutagenesis to enhance the kinetic properties of the involved enzymes. Hence, they constitute highly time intensive, manual efforts with the goal of subsequent implementation of specific pathways in an *in vitro* setting, while computational tools play a minor role in the design and optimization process.

However, such recombinations of pathways can be further explored by computational approaches. Previous work on computational exploration of the carbon fixation space [20] focuses on an in-depth analysis of available data from databases with manual curation, flux balance analysis, and activity analysis for known artificial and natural pathways. The proposed modified pathways are then manually curated following expert intuition [20]. This approach requires complete knowledge of the parameters for every involved enzyme, making generative experiments infeasible.

In this paper, we focus on the computational angle of pathway design itself, for which we propose an approach based on generative chemical space expansion, pathway queries, and topological optimization of pathway solutions based on thermodynamic annotation. The overall goal is fast and flexible pathway suggestions for speeding up the design process. In detail, we present a graph-based computational approach using the cheminformatics software MØD [21], which includes facilities for pathway search via Integer Linear Programming (ILP) optimization [22]. This method allows us to systematically construct and explore chemical spaces with high flexibility, enabling targeted queries for user defined questions, like finding autocatalytic cycles and searching for alternative products. A graphical overview of the approach is presented in Figure 1.

**Figure 1.**
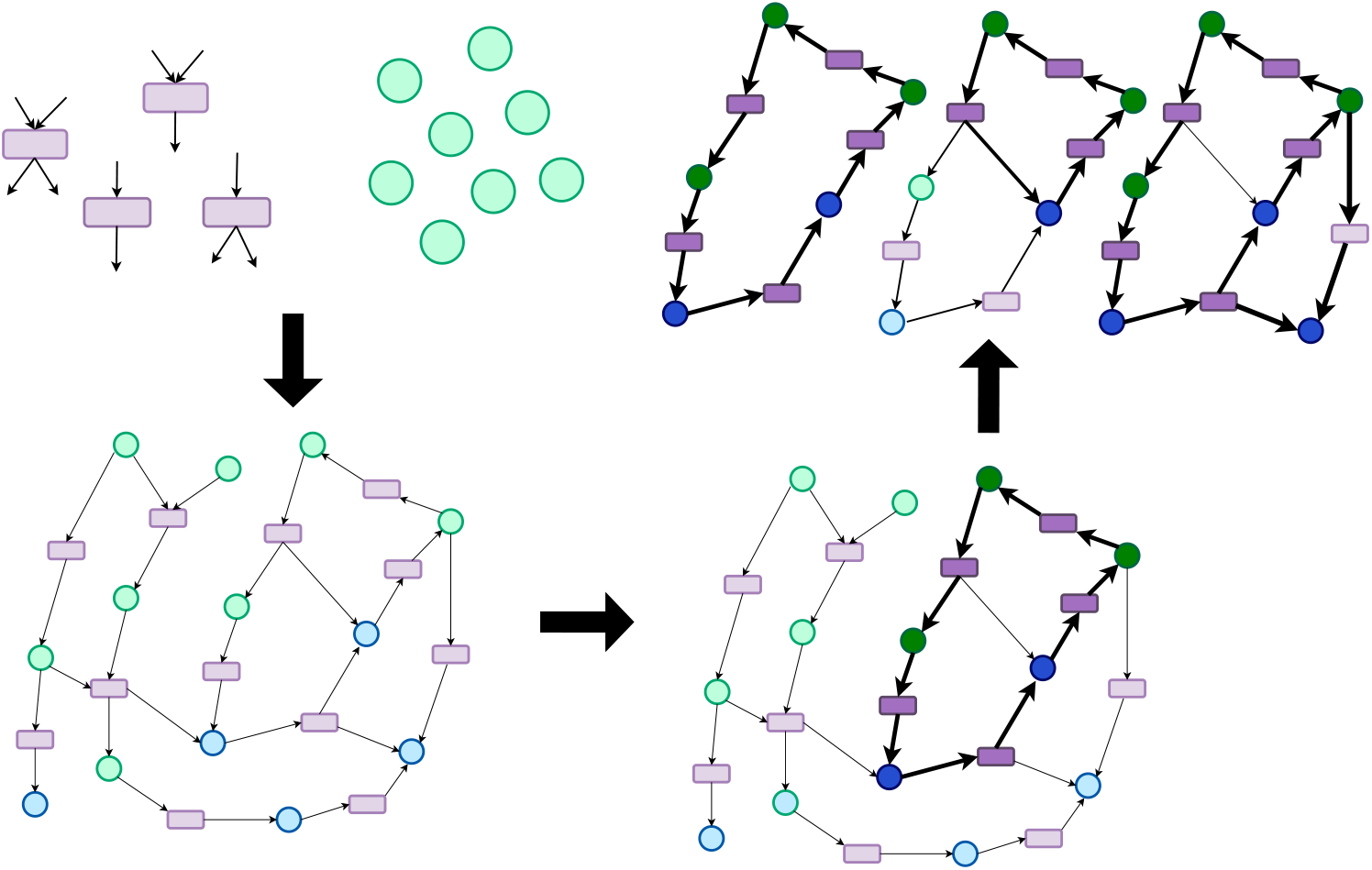
Graphical abstract of the approach for pathway design applied in this study. Graph-grammatical building blocks like rules (in purple) and starting molecules (in green) are used to expand a chemical space. During the expansion, new molecules are created (in blue). On this space, an ILP search is performed to find pathways, where the solutions are evaluated. Highlighted in darker colors are the different pathway solutions.

Our approach starts by iteratively expanding a chemical space, specified by a set of reaction rules and a pool of initial molecules, into a chemical reaction network (CRN). The CRN is constructed as a directed hypergraph, wherein the hyperedges are the reactions and the vertices are the molecules. Reactions are described as reaction rules, each rule representing a general molecular transformation—i.e., a reaction class—instead of a specific reaction (see Figure 5 for an example). This allows the model to capture enzyme promiscuity and account for unconventional or novel metabolic transformations.

Included in the set of rules used in the present study are rules derived from the reactions in the autocatalytic carbon fixation cycles of the Acetyl-CoA-Succinyl-CoA pathway family [3] and in a selection of artificial cycles [18, 23]. The information for the enzymatic reactions involved was taken from the KEGG database [24]. A detailed description of the chemical space composition can be found in the Methods section 4.1.

The initial pool of molecules used to start the stepwise expansion of the CRN consists of known metabolites and common cofactors typically present in the organisms of interest. As the reaction rules are applied to the input set of molecules, they generate new molecules and reactions, progressively expanding the chemical space. Novel molecules will form through the application of promiscuous reaction rules on the input molecules, achieving the expanded chemical space of carbon fixation, represented as a hypergraph, after a set number of expansion steps.

On this directed hypergraph, the search for pathways is achieved through hyperflow queries [22], which are described in detail in Section 4.2.1. In short, a hyperflow is a route through a hypergraph, in this case through a CRN, with an inflow and outflow of some molecules. The net reaction represented by the hyperflow is the flow on the input and the output molecules. The flow on molecules throughout the rest of the route balances out, i.e., each molecule is produced and consumed in equal amounts. Thus, a hyperflow corresponds well with the standard notion of a pathway. A hyperflow query is the search for such a route with a specified structure. The search query can for instance require certain input and output molecules as start and/or end points, as well as forbidden or preferred reactions or elements of the route. These specifications are fed into an ILP model, which is a set of linear equations, with an objective function to be solved and constraints to be satisfied that combined model the desired hyperflow structure. In our work, we use the hyperflow queries to find the shortest carbon fixation pathways in the generated CRN. Specifically, our queries ask for flows minimizing both the overall number of hyperedges (reactions) used and the absolute flow on these respective edges. Since we are interested in autocatalytic pathways, the only allowed inflow and outflow for the model is the autocatalyst itself and cofactors. This way, the net reaction for a solution to the query represents the production of one more autocatalytic molecule under the usage of cofactors.

The feasibility of the solutions found is evaluated by a post-annotation workflow in three ways. First, the length of a pathway solution is compared to pathways from the literature. Second, the number of cofactors is counted as a measure for energy units and electrons used in the pathway. Third, the Δ_*r*_*G*^*′*^ of the reaction is calculated as a measure for the thermodynamic feasibility. This reaction energy Δ_*r*_*G*^*′*^ is calculated by subtracting the combined energies of formation of the educts from the products, with the energies of formation being obtained from using the eQuilbrator API [25, 26]. The workflow is summarized in Figure 1 and a detailed description of the workflow can be found in Section 4.

In Section 2, we give the details of our results. They can be summarized as follows: We suggest novel synthetic carbon fixation pathways, found through recombining elements of the biochemical pathways of the carbon fixation space, that have qualities similar to those of the natural pathways and of some of the most effective synthetic pathways. In addition, we demonstrate the computational efficiency of our methodology by providing 1000 solutions to a variety of pathway queries. It is worth noting that although we illustrate our workflow as an approach for carbon fixation pathway design, applying it to the chemical space of natural and artificial carbon fixation, the generality of the methodology allows it to be used for pathway investigations on any chemical space of interest.

## 2 Results

### 2.1 Chemical Space Design

To expand the chemical space of carbon fixation into a CRN, input molecules were defined and reaction rules were applied to those molecules. The input molecules consist of 49 general molecules that are intermediates of the Acetyl-CoA–Succinyl-CoA pathways [4], the synthetic CETCH cycle by Schwander et al. [18], and the theoretical glyoxylate cycle cycle proposed by Bar-Even et al. [23], and 20 helper molecules, which include molecules like various cofactors, water, and CO_2_. A detailed listing of the input molecules can be found in the graph grammar files in the GitHub repository https://github.com/anne-susann/C_fixation_pathway_design.

The smallest possible CRN in our setup contains the input rules of the carbon fixation cycles as described above. This space was created via one expansion step, leading to 165 vertices, corresponding to molecules and 220 hyperedges, corresponding to reactions, as described in Table 1. In the following, vertices will be referred to as molecules and hyperedges as reactions, for ease of exposition.

**Table 1.** Characterization of the CRN for Carbon Fixation after different expansion rounds. The number of expansion steps represents how often the rules were applied on the chemical space, and the numbers of vertices (molecules) and hyperedges (reactions) describe the size of the CRN.

| Expansion Steps | # Vertices (Molecules) | # Hyperedges (Reactions) |
| --- | --- | --- |
| 1 | 165 | 220 |
| 2 | 318 | 942 |
| 5 | 996 | 29266 |

Performing flow queries, i.e., the search for an autocatalytic carbon fixation pathway through the reaction network as described in Sections 1 and in full detail in 4.2.1, found only the pathways given as input. No recombinations (i.e., no cross connections between the natural and/or synthetic and theoretical pathways) within the chemical space took place with only one expansion step.

With two expansion steps, the number of reactions increases to 942, while the number of molecules nearly doubles to 318 (see Table 1). This is the space that subsequent analyses were performed on. This CRN proved big enough to have novel compounds and novel cross connections and pathways, but does not reach a size where running flow queries requires a large amount of computation time.

We experimented with further expansion steps up to and including five, at which point combinatorial growth rendered flow queries computationally infeasible. While larger expansions were technically possible, they resulted in disproportionately large networks with limited additional insight. Earlier steps (three and four expansion steps) were explored, but the two-step expansion space already supported sufficient pathway choices and kept computation times for flow queries down.

### 2.2 Pathways Compared to Literature

Various comparison studies of autocatalytic carbon fixation cycles have introduced the concept of comparing a set of key measures [4, 18, 27] to understand the classification of a novel pathway. These measures include the number of steps required to complete one cycle, the ATP units and cofactors used in the completion of this cycle, and the carbon units fixed per cycle.

In this work, we propose two novel theoretical pathways found via our methodology. The two presented pathways are the result of flow queries searching in the CRN for the shortest pathway (that is, with the fewest number of reactions) adhering to the structural definition of autocatalytic cycles in Section 1. The autocatalyst, i.e., the inflow and outflow molecule, in the two searches were Acetyl-CoA and Malate, respectively, and the query specified the production of one additional molecule of the autocatalyst while fixing carbon in the form of CO_2_ or HCO_3_^2 –^. This search was performed on the CRN obtained after two expansion steps and characterized in the second line of Table 1. The two proposed theoretical pathways we refer to according to their search objectives as Shortest autocatalytic cycle Acetyl-CoA, and Shortest autocatalytic cycle Malate. In Table 2, we compare their characteristics to benchmark pathways from other studies. The explicit structure can be found in the GitHub repository under Output Pathways https://github.com/anne-susann/C_fixation_pathway_design.

**Table 2.**
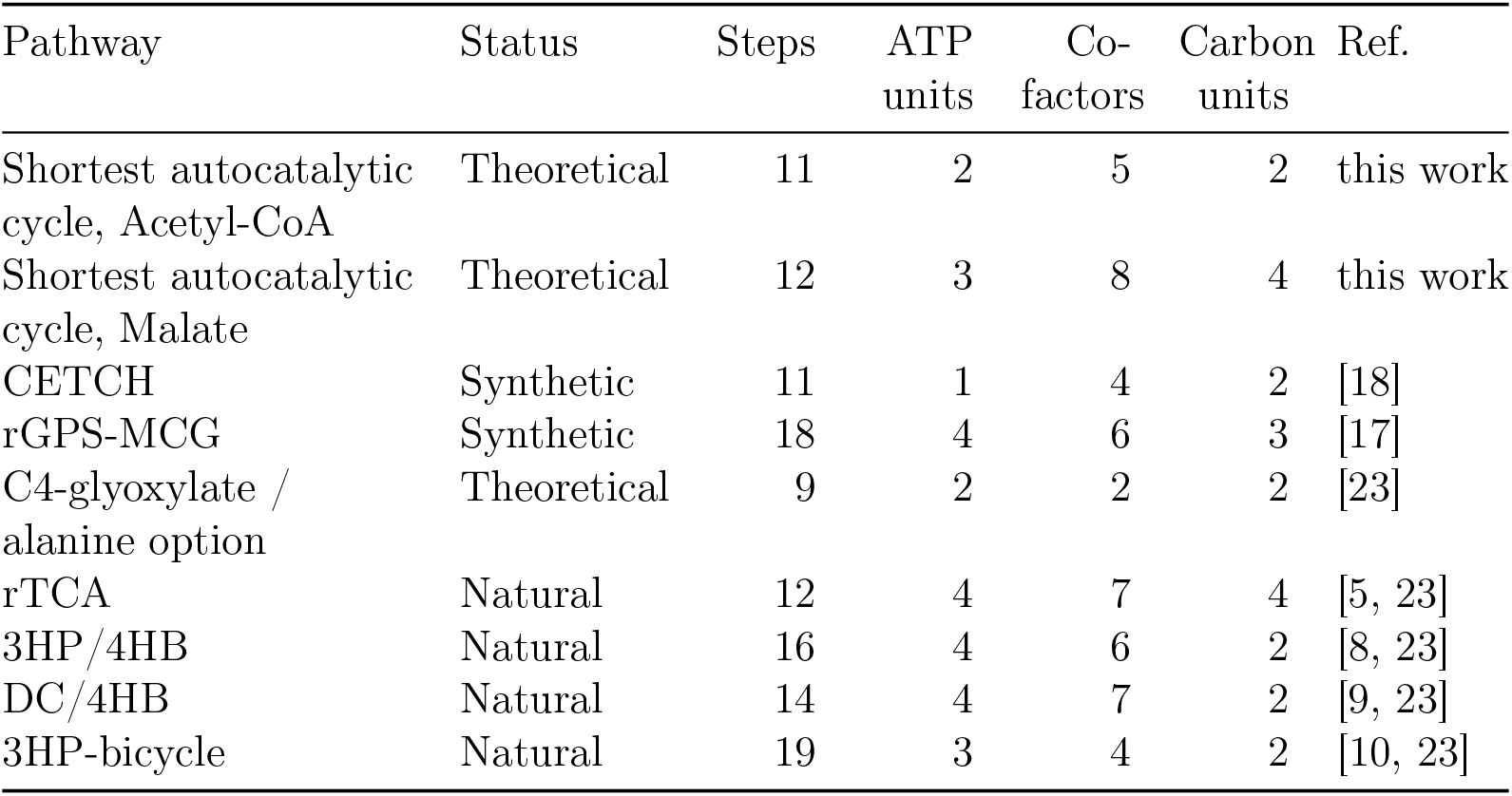
A comparison of the flow solutions as proposed theoretical carbon fixation pathways with selected theoretical, synthetic, and natural pathways. Pathway: name of the autocatalytic carbon fixation cycle. Status: implementation status of a given pathway, divided into the categories theoretical, implemented as synthetic, or known natural pathway. Steps: number of steps necessary to complete one autocatalytic cycle and produce one additional molecule of the autocatalyst. ATP units / Cofactors / Carbon units: number of units or molecules needed for one autocatalytic cycle to complete. Ref.: Reference paper where the pathway was discovered or described. CoA: Coenzyme A, CETCH: Crotonyl-CoA/ethylmalonyl-CoA/hydroxybutyryl-CoA, rGPS-MCG: reductive Glyoxylate Pyruvate synthesis Malyl-CoA-glycerate, rTCA: reductive tricarboxylic acid, 3HP/4HB: 3-hydroxypropionate 4-hydroxybutyrate, DC/4HB: dicarboxylate

These benchmark pathways are either natural, theoretical, or synthetic, the last phrase meaning that they have been theoretically designed and then implemented *in vitro*. The Acetyl-CoA pathway requires 11 steps to generate one autocatalytic molecule, and the Malate pathway takes 12 steps. Compared to the natural pathways, the two novel suggestions either use strictly fewer or the same number of steps as the shortest natural one. Compared to the synthetic pathways, the novel suggestions are in the range of the shortest synthetic pathway in terms of number of steps used. ATP units and cofactors as energy and reduction equivalents give a cost measure for cells or in vitro systems. The Acetyl-CoA pathway requires one unit more of ATP (2 in total) and one more reduction-oxidation (redox) cofactor (5 in total) than the highly efficient CETCH cycle while fixing the same amount of carbon units per cycle, namely two. It is comparable in ATP and cofactor usage to natural carbon fixation cycles when normalizing for carbon units. The Malate pathway has a higher total requirement, with 3 ATP units and 8 redox cofactors, but fixing 4 carbon units in the process, making it more efficient than the Acetyl-CoA cycle. The Malate cycle is also on par with the shortest natural pathway, rTCA, with respect to these cofactor requirements.

### 2.3 Exploration Scope

The exploratory potential of our approach includes the search for many good solutions, not just a single best. Three different queries for autocatalysis, using Acetyl-CoA, Malate, and Propionyl-CoA as autocatalytic molecules respectively, were taken as benchmark searches for investigating the differences in the solutions for different biological products. The flow query was built to find 1000 solutions satisfying the query constraints. Within those, as long as the objective value (that is, pathway length) is the same, the solutions are considered equally good by the ILP solver. However, each of the 1000 solutions were required to be topologically different. These 1000 solutions were then evaluated statistically by the number of cofactors that are used in each solution and by the reaction energy Δ_*r*_*G*^*′°*^ value of those solutions, visualized in Figure 2.

**Figure 2.**
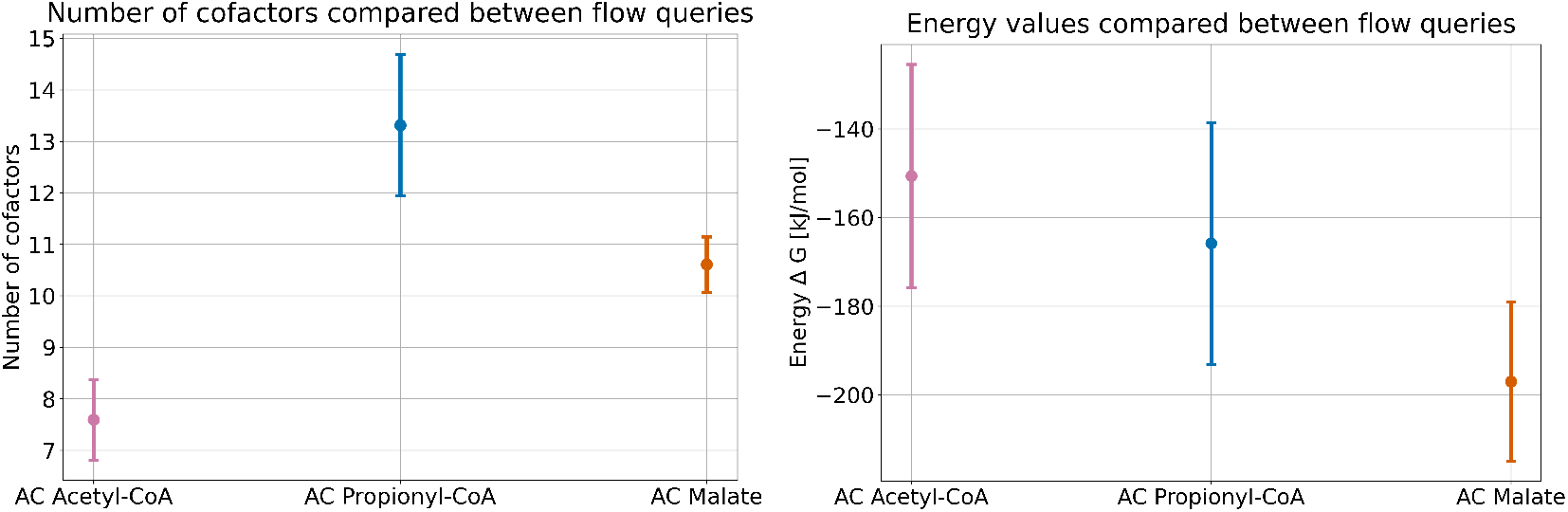
Comparison of cofactor usage and reaction energy for 1000 solutions to different flow queries. The cofactors include ATP units and redox cofactors such as NAD(P)H and Ferredoxin. The three queries are looking for the shortest autocatalytic cycle to produce Acetyl-CoA, Propionyl-CoA, and Malate, respectively. Represented is the average with the bars indicating the minimum and maximum outliers.

The two measures shed light on two different parameters for a potential implementation, namely cofactor requirement and thermodynamic feasibility. A more negative Δ_*r*_*G*^*′°*^ makes a pathway in general more likely to react in the forward direction, while the cofactor requirements are generally considered the standard measure of energy efficency in systems biology. While a negative value for Δ_*r*_*G*^*′°*^ is required, the amount of cofactors used in a pathway heavily influences its implementation potential, since cofactor-hungry pathways require more input of expensive cofactors and the addition of recycling pathways to the setup.

The results show that the Acetyl-CoA pathway has the lowest usage of cofactors, with an average of 7.6 cofactors used (see Figure 2 and Table 3). The Propionyl-CoA and Malate pathways need more cofactors, with an average of 13.3 and 10.6. Since the latter two pathways are longer in steps, the cofactors were also calculated per step of the pathway. This results in a cofactor-per-step ratio of 0.69 for Acetyl-CoA, and 0.89 and 0.88 for Propionyl-CoA and Malate (see Table 3), making the autocatalytic Acetyl-CoA cycle not only the shortest but also the most cofactor-efficient pathway.

**Table 3.** Flow query results for the 1000 solutions to find the shortest autocatalytic carbon fixation pathway, for Acetyl-CoA, Malate, and Propionyl-CoA, respectively. The table shows the autocatalytic molecule that was searched for, the average length of the pathway solution as steps, the average co-factor use and the co-factor use normalized against the pathway length.

| Autocat. molecule | Steps average | Co-factors average | Co-factors per step | Energy average [kJ mol<sup>-1</sup>] |
| --- | --- | --- | --- | --- |
| Acetyl-CoA | 11 | 7.6 | 0.69 | -150.66 |
| Propionyl-CoA | 15 | 13.3 | 0.89 | -165.82 |
| Malate | 12 | 10.6 | 0.88 | -196.98 |

The energy measure shows Malate having the lowest energy value for the pathway, with Δ_*r*_*G*^*′°*^ = *™*196 kJ mol^*™*1^ on average. The Acetyl-CoA pathway has the highest energy with Δ_*r*_*G*^*′°*^ = *™*150 kJ mol^*™*1^, and Propionyl-CoA is in the middle with Δ_*r*_*G*^*′°*^ = *™*165 kJ mol^*™*1^. In this measure, the Malate pathway seems to be the most thermodynamically driven, especially when regarding the need for one more step compared to Acetyl-CoA but still having a lower energy value. However, these Δ_*r*_*G*^*′°*^ values include the energy gained from the used cofactors.

### 2.4 Individual Solutions Show Novel Acetyl-CoA Producing Pathways

As a benchmarking example, three individual solutions from the solution space for shortest autocatalytic Acetyl-CoA production were further inspected. The solutions were chosen based on heuristic considerations on topological comparability from the solution space mentioned before. The topological structure of the three solutions is very similar, hence we can overlay and compare their specific differences in steps. In Figure 3, the common core of the three solutions is the parts where molecules are shown on a white background while the respective differences are marked by green, pink, and orange. Acetyl-CoA as the autocatalytic molecule, marked in yellow, is at the center of the pathways. Much of the pathway relies on the rTCA as the basic structure, but we find a glyoxylate shunt in between, as well as a combination of other reactions of the Acetyl-CoA-Succinyl-CoA space. An important branchpoint between the three solutions is on the route between Oxalacetate and Malyl-CoA, where we observe different combinations of enzymes used and intermediate molecules produced.

**Figure 3.**
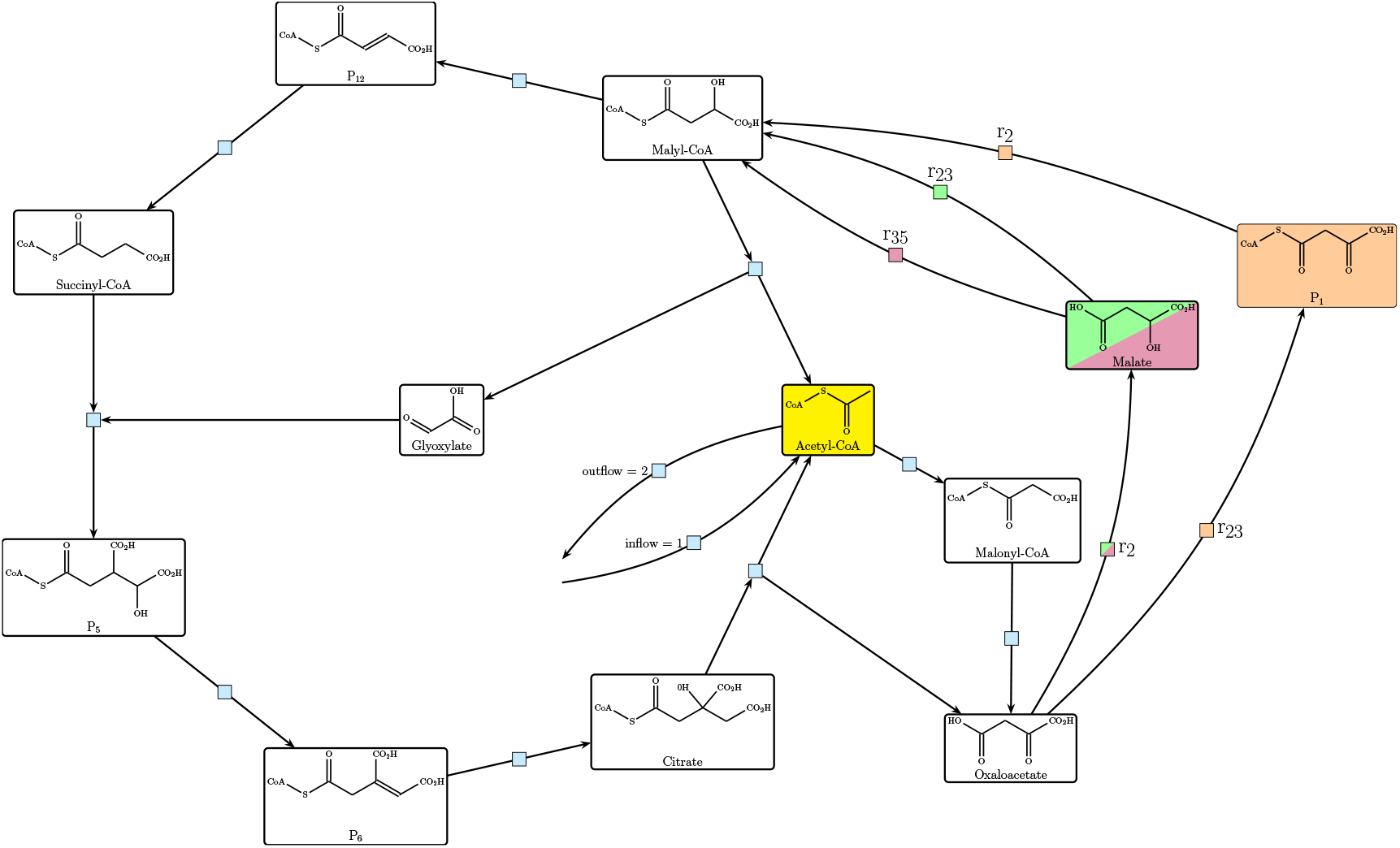
Comparison of the first 3 solutions for finding the shortest autocatalytic cycle producing Acetyl-CoA. The important differences are in the rules used in the solution and the molecules used. Reaction rules used are modeled after the following enzymatic reactions: *r*_16_, Acetyl-CoA-carboxylase (3HP4HB); *r*_9_, 2-ketogluterate:ferredoxin oxidoreductase (rTCA); *r*_2_, ATP citrate lyase (rTCA); *r*_23_, 4HB-CoA synthetase (3HP4HB); *r*_35_, succinyl-CoA malate CoA transferase (3HP bicycle); *r*_20_, 3HP-CoA dehydratase (3HP4HB); *r*_5_, fumarate reductase (rTCA); *r*_54_; *r*_6_, fumarase (rTCA); *r*_7_, aconitase (rTCA); *r*_8_ crotonase (rTCA); *r*_1_, succinyl-CoA synthetase (rTCA).

Because of the difference in enzymes used between the compared solutions, the cofactor usage also changes. In Table 4, the inflow and outflow of cofactors for each solution is detailed. The main difference between the pathways is the ATP usage, with solution 1 (Sol 1 in Table 4) having the lowest ATP consumption of only one. This yielded a higher 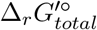 value, but with *™*80 kJ mol^*™*1^ the pathway is still thermodynamically feasible and has a low ATP usage.

**Table 4.**
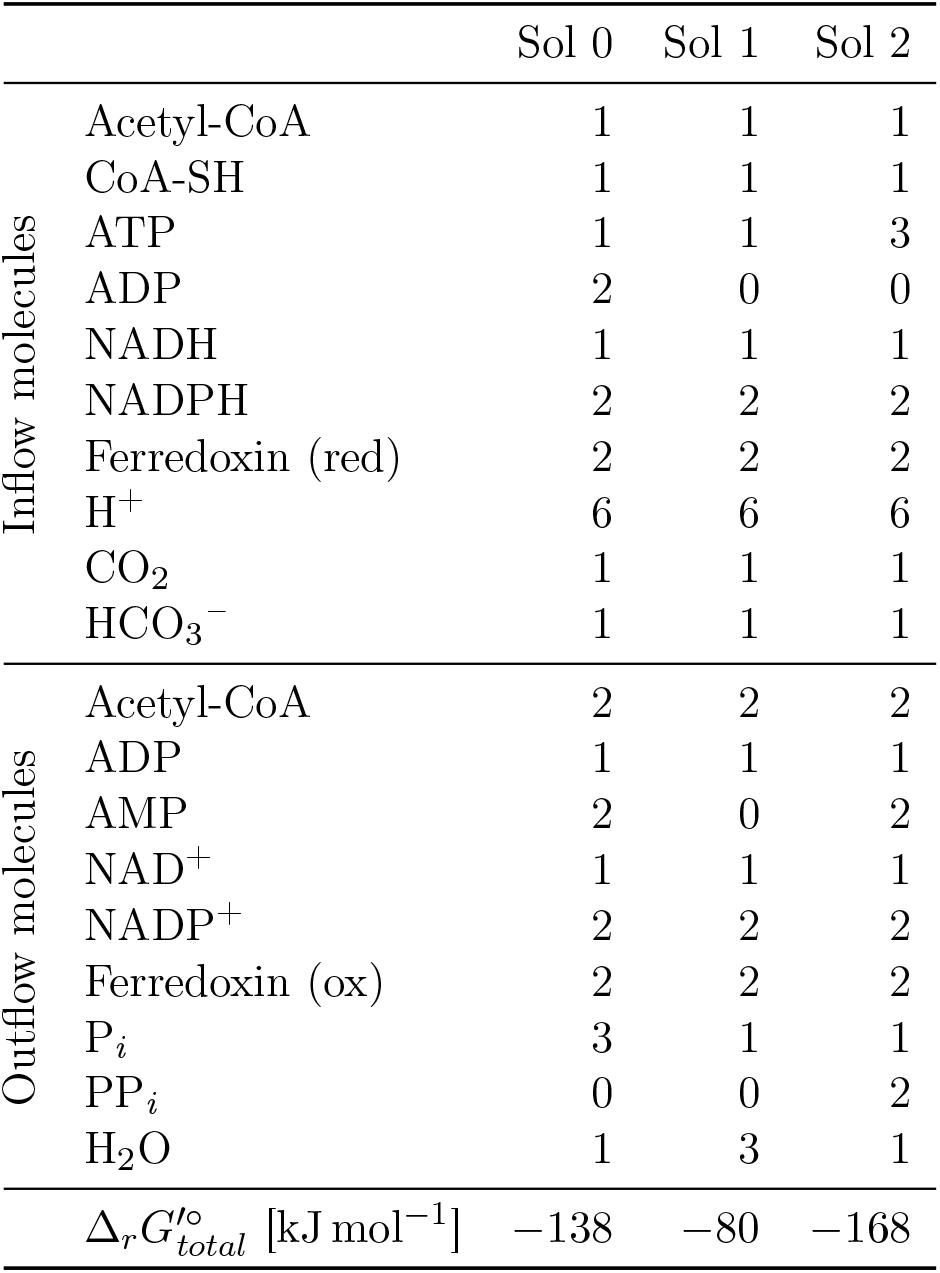
Detailed description of the inflow and outflow of molecules as well as the overall reaction energy for the three flow solutions depicted in Figure 3. The flow solutions were optimized to find the shortest autocatalytic pathway to produce Acetyl-CoA in the carbon fixation space.

A closer view on the three flow solutions shows the combination of natural pathways to form novel pathways, as on the topological level, we find a reordering of enzymatic reactions and recombination of reaction rules with different natural pathway origins.

## 3 Discussion

In this paper, we present a new method to explore the chemical space of carbon fixation for novel pathway combinations as well as artificial pathways, and we give examples of novel pathways. In our experiments, the space was restricted to be based on known natural, synthetic, and theoretical carbon fixation pathways, but the method can be generalized to any chemical space, natural or not, involving any number and types of reactions. The chemical space expansion strategy allows for a flexible design of the CRN and an exploratory approach to investigate the chemical space.

We found two interesting novel pathways after applying our search framework, one producing Acetyl-CoA and the other producing Malate. Both autocatalytic cycles have quality measures comparable to other artifcially designed pathways and the most efficient natural pathways, as detailed in Section 2.2.

The lower energy value of the Malate pathway makes it an interesting contender for potential implementation. However, we note that the overall cofactor usage of the Acetyl-CoA pathway as described in Figure 2 and Table 3 is lower while the pathway still has a prominent negative Δ_*r*_*G*^*′°*^ as a driving force. In a wet-lab implementation setting, this means that less cofactors need to be supplied and recycled by a cofator recycling system. This likely makes the Acetyl-CoA pathways the more interesting platform for further exploration.

Another factor to be considered with respect to feasibility of implementation is the specific cofactors used in the solutions. The pyruvate synthase (pyruvate:ferredoxin oxidoreductase) and the ketoglutarate synthase (2-ketogluterate:ferredoxin oxidoreductase) are both highly efficient carboxylating enzymes found in the rTCA. They use Ferredoxin as the reduction cofactor, which has a higher reduction potential than NAD(P) [3]. We can see in Table 4 that the solutions compared here all apply Ferredoxin, as do the solutions explored in Section 2.3. However, Ferredoxin is an oxygen-sensitive cofactor, meaning implementation would have to be under anaerobic conditions. This could be a major hurdle for the in vitro implementation of the suggested pathways, but it could possible be achieve through metabolic engineering of anaerobic strains [28].

A deeper look into the composition of the Acetyl-CoA pathway solutions revealed that the combination of enzymes from different natural origins yields shorter and more efficient pathways. The specific combination in each solution can differ. At the same time, the length of the pathway stays the same, while the cofactor usage differs. This shows the versatility of the overall chemical space and highlights the potential to get many pathway suggestions, which can then be filtered to satisfy relevant criteria for later implementation, such as oxygen-sensitivity, ATP usage, or redox cofactor usage.

We were able to achieve these results while using a strictly topological approach for pathway searches, with ILP as the search algorithm for novel pathways. This approach, augmented with a post-annotation workflow for feasibility measures, gives a flexible and robust approach for exploratory pathway searches in any chemical space. At the same time, there is a minimal need for extensive database searches and the associated computational cost is low, as all of the workflow can be performed on a standard laptop.

The advantage of our approach is the possibility to individualize the optimization criteria, to include as many constraints as wanted, and to explore beyond the scope of what is known already. By the rule based expansion of the CRN, it can produce molecules previously unknown to a specified chemical space and can apply the given reactions to those molecules as well, while still adhering to the chemical principles of a reaction network [29]. Future work should consider kinetic measures as well as focus on including cofactor recycling systems, in order to make the proposed solution less cofactor expensive and more promising to implement in a wet lab setting. Future design efforts may also focus on incorporating metabolic reactions not directly involved in carbon fixation pathways to look for promising short cuts.

## 4 Methods

The exploration of the carbon fixation space and the search for artificial pathways followed the workflow shown in Figure 4. The workflow can be split into the following tasks: (i) selection of the chemical space that will be modeled, (ii) expansion of the chemical space using MØD, (iii) optimization and pathway search, and (iv) post annotation of pathway solutions. Each of these tasks will be explained in detail in the following.

**Figure 4.**
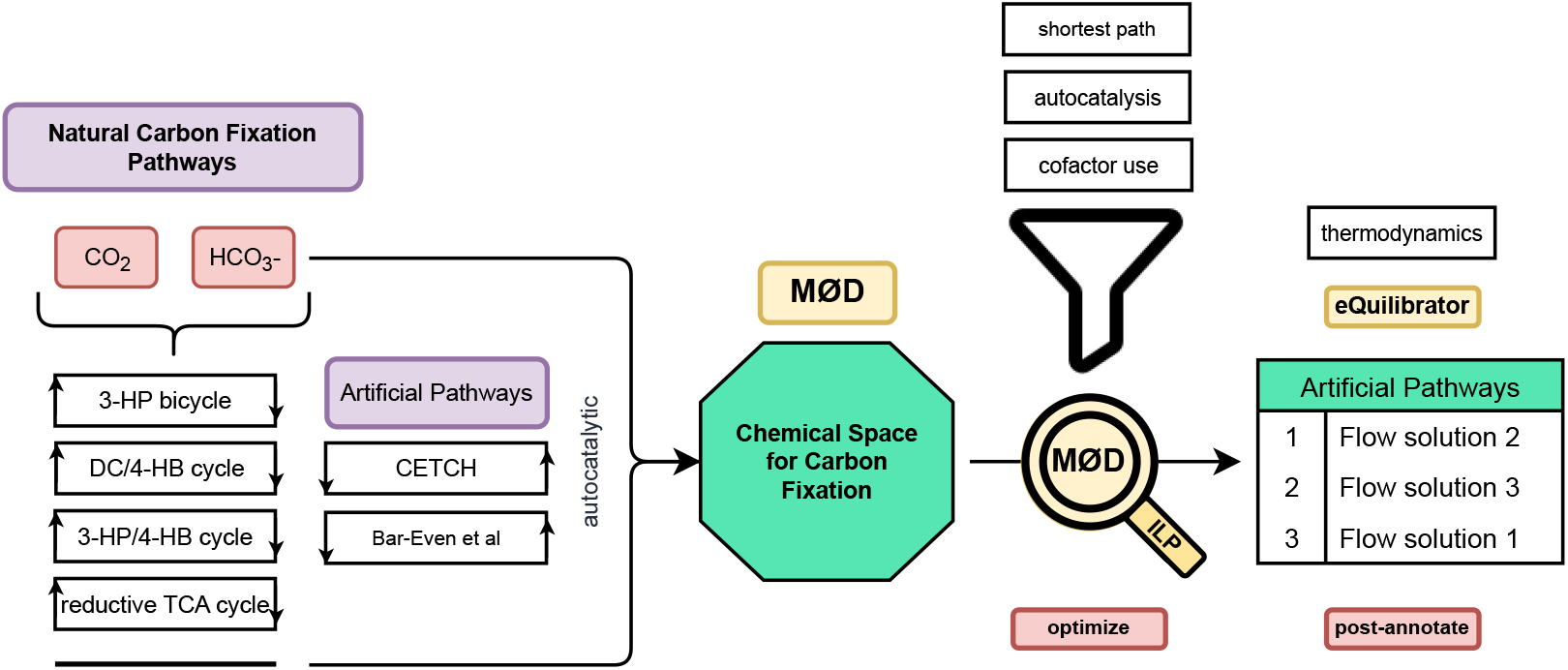
Workflow for using a graph-based rule-based approach for the exploration of the chemical space of natural and artificial carbon fixation. First, the natural and artificial pathways are turned into transformation rules and the chemical space is expanded. Then, flow queries are used to search and optimize for a given objective function with constraints, yielding a list of flow solutions. Lastly, post-processing allows for ranking of solutions. 3-HP: 3-hydroxypropionate, DC: dicarboxylate, 4-HB: 4-hydroxybutyrate, TCA: tricarboxylic acid cycle, CETCH: crotonyl-CoA/ethylmalonyl-CoA/hydroxybutyryl-CoA cycle [18], Bar-Even et al.: proposed pathways from Bar-Even and his colleagues [23]

### 4.1 Modeling the Chemical Space of Carbon Fixation

The chemical space of carbon fixation was formalized using MØD [21], a graph- and rule-based cheminformatics tool capable of constructing chemical reaction networks as directed hypergraphs. A detailed description of the software MØD, including the hypergraph construction, can be found in Andersen et al. [21, 30, 31].

The hypergraph in our approach is the representation of the CRN. A CRN generally consists of a set of molecules and a set of reactions. This can be modeled as a directed multi-hypergraph *H* = (*V, E*), with the set of vertices *V* representing the molecules, and the set of directed hyperedges *E* representing the reactions. Each hyperedge *e ∈ E* contains a pair (*e*_*tail*_, *e*_*head*_) of multisets of vertices *e*_*tail*_ *⊆ V* and *e*_*head*_ *⊆ V*, corresponding to the molecules that flow in and out of a reaction. Hyperedges, as opposed to simple edges, allow the direct modeling of many-to-many relations between reactant and product molecules.

CRNs in MØD are expanded using three core elements. Firstly, rules represent the (bio)chemical reactions and define how molecules are transformed. Rules operate as graph transformations that rewrite molecular graphs, breaking or forming bonds and adjusting charges. Depending on the molecular context attached to the reaction center of the rewite rule, the specificity of the rewrite rule may be tuned. In this way one rule may represent multiple reactions that induce the same net change in the molecules it may be applied to. For example, a generic reduction rule with no context around the reaction center will perform all reactions catalyzed by oxidoredutases like reductases or dehydrogenases (see Figure 5). This allows for a flexible construction of the chemical space, not needing to formalize every single reaction but working with reaction types and classes.

**Figure 5.**
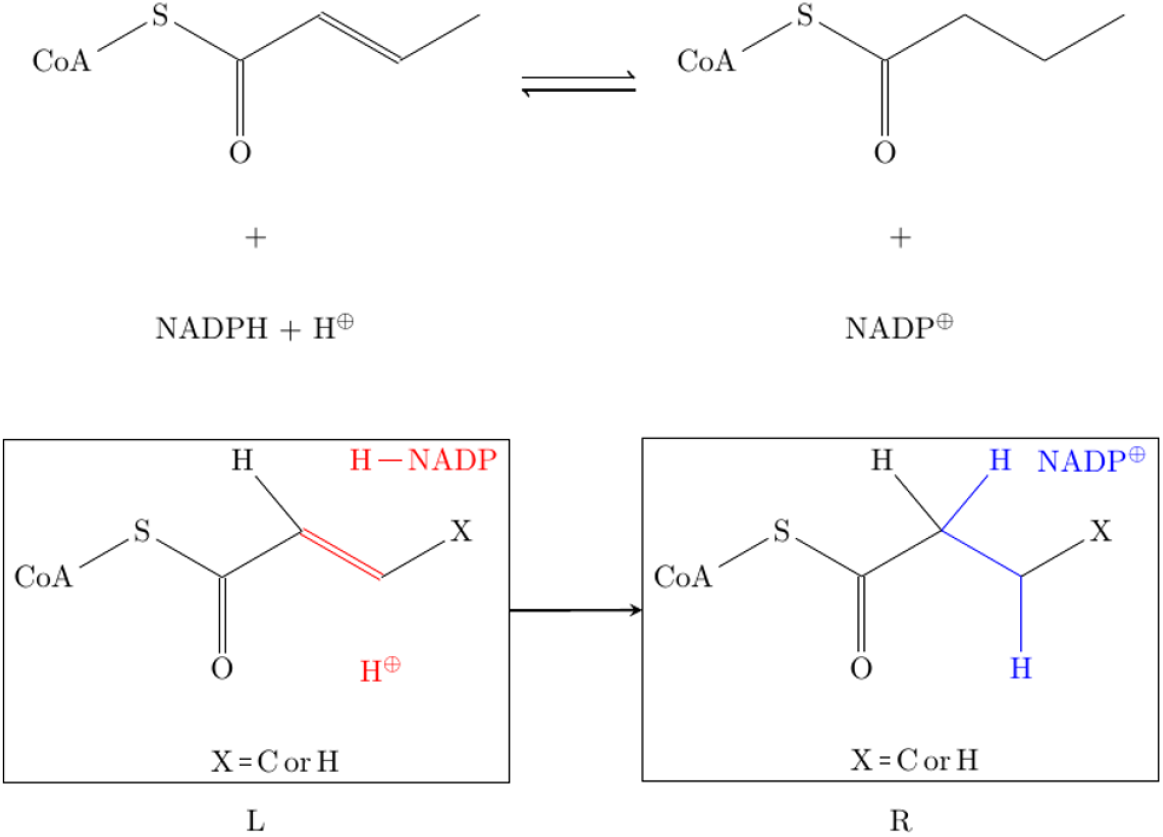
Example of a transformation rule as used by MØD. This specific rule shows the acrylyl-CoA reductase as found in the 3HP/4HB cycle of carbon fixation. The mechanism was modeled after the respective KEGG entry for this enzyme [24]. The top shows a traditional representation of the enzymatic reaction, while the bottom shows the representation as a graph transformation rule, with the left (L) and the right (R) side as the bonds before and after the reaction. X is a variable that can define different atoms, in this case C or H.

Secondly, molecules serve as the input graphs upon which the rules are applied, expanding the network from a set of core molecules. Finally, expansion strategies define how the rules are applied iteratively to the given molecules to construct the reaction network step-by-step. A more detailed description of CRN expansion in MØD can be found here [30]. For the networks built here we used an expansion strategy that restricts the chemical space to molecules with at most 6 C atoms, to avoid a combinatorial explosion and stay close to nature in this approach. All of the included natural pathways and the known artificial pathways have no molecules involved that go beyond 6 C atoms in the backbone of the molecule. The carbon atoms in the covalently bound cofactor Coenzyme A (CoA) are not considered for this 6 C restriction. Further, the expansion was restricted to produce molecules with at most one CoA attached to them, to maintain biochemical validity and stability in the CRN.

The rules were derived from enzymatic reactions available in the KEGG database [24]. For this study, the molecular context for the reactions were designed conservatively to reflect enzymatic specificity while allowing for some flexibility that could reasonably be reached with enzyme isoforms, enzyme engineering or enzyme promiscuity. This study focused on autocatalytic carbon fixation cycles, excluding pathways like the Wood-Ljungdahl pathway and the reductive glycine pathway that are not autocatalytic [14, 16]. The Acetyl-CoA–Succinyl-CoA pathways (e.g., rTCA, DC/4-HB, 3-HP/4-HB, 3-HP bicycle) were central to the model due to their inherent autocatalysis and overlapping structures. The synthetic CETCH cycle [18], which builds on these natural pathways, was also included, as well as proposed pathways from a paper by Bar-Even and his colleagues [23]. The Calvin-Benson-Basham (CBB) cycle [6], despite being autocatalytic, was not included in the explored space due to its unique molecular context, which relies on recombination and carbohydrate chemistry, and lack of interaction with other cycles.

### 4.2 Finding Novel Artificial Pathways

#### 4.2.1 The ILP Model

To identify novel pathways within the modeled chemical space, we utilized integer linear programming (ILP), implemented within MØD as “flow queries” [22, 16]. In this context, a pathway is seen as a hyperflow on the chemical reaction network, and a pathway query is therefore a flow query, with constraints on the flow, as visualized in Figure 6.

**Figure 6.**
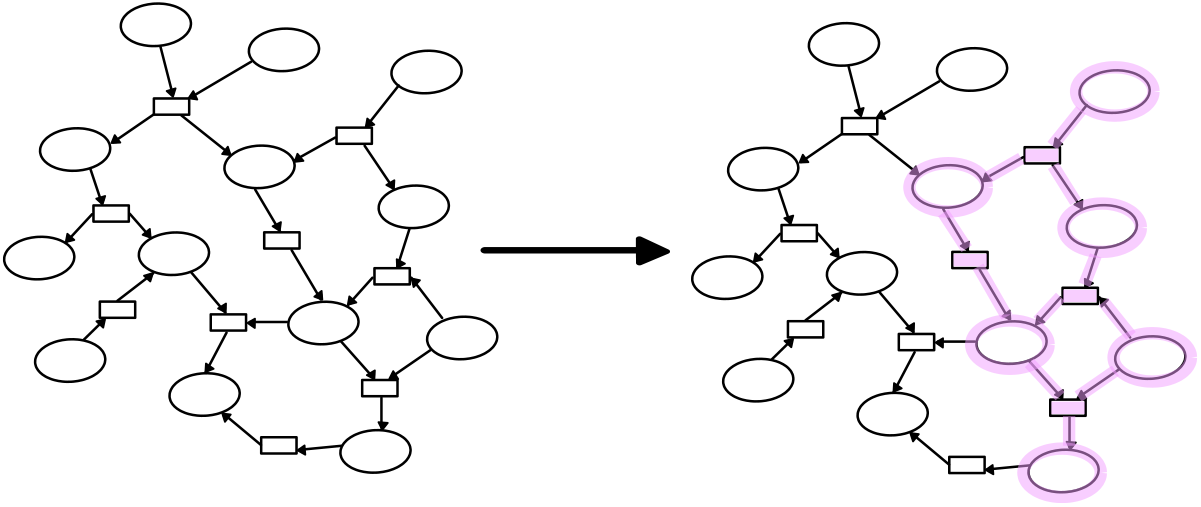
A flow query is a pathway search on a chemical reaction network. On the left is the network, with ovals representing molecules and rectangles representing reactions. The flow query is a set of constraints and an objective function which combined specifies desired structural aspects of the pathway searched for. A solution to a flow query is illustrated on the right.

These searches are built with objective functions and constraints as described below. The integer nature of the ILP model allows for integer values as flow solutions, which therefore represent only whole molecules in the solution space. Unlike classical flux balance analysis, which emphasizes continuous flux distributions under steady-state conditions [32], the ILP approach with integer solution values focuses on the structure of a pathway, identifying which reactions are active rather than optimizing the flux magnitude (see [22] for a detailed comparison).

Defining the objective function focuses on the main driving force of this work, the optimization towards shorter artificial pathways. The main objective function is described in Equation (1). A description of the appearing variables and constants is given in Table 5. Shortest pathway searches can be achieved by minimizing the total use of edges in a solution, represented by *z*_*e*_. Additionally, to avoid scaled solutions with same edge numbers but up-scaled flow variable, *x*_*e*_ is minimized as well.

**Table 5.** Variables and constants used in the ILP formulation.

| Symbol | Type | Description |
| --- | --- | --- |
| $z_e$ | boolean variable | Indicator variable for flow on hyperedge $e$ |
| $x_e$ | integer variable | Flow variable for a hyperedge $e$ |
| $w$ | constant | Weight to prioritize minimized edge usage, set to 1000 |

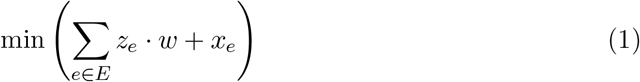

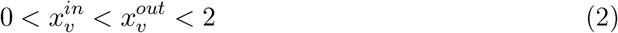

To find a hyperflow in the network, a set of sources and sinks needs to be defined. The sources are the molecules that we allow to have inflow, i.e., the molecules that can be put into the pathway. The sinks are the molecules that we allow to have outflow, i.e., the molecules that can be eventually produced by a pathway. In order to find an autocatalytic flow solution, only the autocatalytic molecule and the cofactors can be allowed as sources and sinks. In this way, the net reaction of the flow has only those molecules consumed and produced. The constraint in Equation (2) assures that the autocatalytic molecule *a* of the query has an input greater than zero and an output greater than the input, thereby defining the net production of at least one autocatalyst molecule. This modeling of autocatalysis follows the definition of autocatalysis presented in [22, 16].

The queries were formulated with objective functions tailored to specific optimization tasks, such as identifying the shortest autocatalytic cycles or finding alternative products. Flow queries were solved using the Gurobi optimizer [33] under an academic license. All computations were performed on a standard consumer laptop (AMD Ryzen 7 5700U, 16 GB RAM, Windows 11). The runtime for the search for 1000 solutions added up to just under 18 hours.

The flow queries were performed on three molecules to represent the customizability of the search. Acetyl-CoA was chosen as a natural choice for a benchmark molecule because all of the included natural pathways have the overlap of this molecule in their pathways, while it also plays a role in many theoretical pathways [23, 17, 3]. Malate is a intermediate molecule in all of the included natural pathways as well as used as a product molecule for the CETCH cycle [18]. Propionyl-CoA was used as molecule that is only found in two of the natural pathways and a common benchmarking molecule in synthetic pathway design [18, 19].

#### 4.2.2 Evaluation of Pathway Solutions

Initially, pathway solutions were manually inspected to identify interesting combinations and validate the presence of known pathways. This step involved comparing key attributes of the proposed pathway. One of these key attributes is cofactor usage, which already gives a measure for energy consumption of a pathway and alignment with natural or proposed artificial pathways. Another one is the number of reaction steps, comparing the pathways length and efficiency in absolute length. Lastly, the involved reactions themselves were compared, determining how closely related a solution is to a known pathway in topology and what recombinations were proposed in the solution. This includes alternative reaction routes, as well as molecules not found in the natural space being used in the solutions. These manual evaluations provided insights into how each pathway diverges from natural pathways and how it mimics or extends known artificial pathways.

Thermodynamic feasibility was subsequently assessed to complement this qualitative analysis. Energies of formation for all molecules in the chemical space were calculated using the component-contribution method via the eQuilibrator tool [25, 26, 34, 35]. The energies of formation were calculated at cellular conditions Δ_*f*_ *G*^*′°*^, with pH = 7 and ionic strength of 0.1 m. As a measure of feasibility the Gibbs free energy change Δ_*r*_*G*^*′°*^ was calculated by post-annotating each flow solution with energies of formation for the overall reaction of the pathway. The sum of Δ_*f*_ *G*^*′°*^ of the reactants was subtracted from the sum of Δ_*f*_ *G*^*′°*^ of the products, as described in Equation (3). The calculation of the reaction energy provided a metric for the analysis of pathway favorability, and includes the thermodynamic activation cost required to create reactive molecules through ATP use.

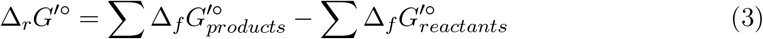

Further insight into the biochemical feasibility was achieved by analyzing the cofactor usage. ATP and ADP were used as a measure for energy usage. The consumption of reduced redox cofactors, including NAD(P)H, Ubiquitin, and Ferredoxin, is a measure of the electrons needed for the pathway. Those two cofactor counts combined give an estimate of how expensive a pathway would be for an organism to perform. Those cofactors were counted in a given solution as part of the post-annotation analysis.

This combined approach of qualitative inspection and thermodynamic evaluation offered a framework for identifying and prioritizing both natural-like and novel artificial pathways for further investigation.

## 5 Code and Data Availability

The datasets generated and analyzed during the current study, as well as the underlying code are available in the GitHub repository https://github.com/anne-susann/C_fixation_pathway_design.

## 6 Author Information

### 6.1 Contributions

C.F., R.F., J.L.A, D.M. conceptualised and supervised the study. J.L.A, N.L. and C.F. provided the computational framework, and with A-S.A. they designed the set-up and experiments. A-S.A. implemented and performed the computational experiments, the data analysis, and the writing and editing of the paper. C.F., R.F., J.L.A, D.M. revised, and with N.L. edited the manuscript.

## 7 Acknowledgments

We thank Prof. Annette Taylor (University of South Hampton) for scientific discussions and her input on the manuscript. This project was funded by the European Unions Horizon Europe Doctoral Network programme under the Marie-Skłodowska-Curie grant agreement No 101072930 (TACsy Training Alliance for Computational systems chemistry). The funder played no role in study design, analysis, and interpretation of the data.

## 8 Ethics Declaration

### 8.1 Competing interests

All authors declare no financial or non-financial competing interests.

## Notes

https://github.com/anne-susann/C_fixation_pathway_design

### Competing Interest Statement

The authors have declared no competing interest.

## References

[1] Michael D. Burkart, Nilay Hazari, Cathy L. Tway, and Elizabeth L. Zeitler. Opportunities and Challenges for Catalysis in Carbon Dioxide Utilization. ACS Catalysis, 9(9):7937–7956, September 2019.

[2] Noor Yusuf, Fares Almomani, and Hazim Qiblawey. Catalytic CO2 conversion to C1 value-added products: Review on latest catalytic and process developments. Fuel, 345:128178, August 2023.

[3] Arren Bar-Even, Avi Flamholz, Elad Noor, and Ron Milo. Thermodynamic constraints shape the structure of carbon fixation pathways. Biochimica et Biophysica Acta (BBA) - Bioenergetics, 1817(9):1646–1659, September 2012.

[4] Arren Bar-Even, Elad Noor, and Ron Milo. A survey of carbon fixation pathways through a quantitative lens. Journal of Experimental Botany, 63(6):2325–2342, March 2012.

[5] M C Evans, B B Buchanan, and D I Arnon. A new ferredoxin-dependent carbon reduction cycle in a photosynthetic bacterium. Proceedings of the National Academy of Sciences, 55(4):928–934, April 1966. Publisher: Proceedings of the National Academy of Sciences.

[6] Andrew A. Benson, James A. Bassham, Melvin Calvin, A. G. Hall, H. E. Hirsch, S. Kawaguchi, V. Lynch, and N. E. Tolbert. THE PATH OF CARBON IN PHOTOSYNTHESIS: XV. RIBULOSE AND SEDOHEPTULOSE. Journal of Biological Chemistry, 196(2):703–716, June 1952.

[7] Stephen W. Ragsdale. The Eastern and Western branches of the Wood/Ljungdahl pathway: how the East and West were won. BioFactors, 6(1):3–11, 1997. _eprint: https://onlinelibrary.wiley.com/doi/pdf/10.1002/biof.5520060102.

[8] Ivan A. Berg, Daniel Kockelkorn, Wolfgang Buckel, and Georg Fuchs. A 3-hydroxypropionate/4-hydroxybutyrate autotrophic carbon dioxide assimilation pathway in Archaea. Science (New York, N.Y.), 318(5857):1782–1786, December 2007.

[9] Harald Huber, Martin Gallenberger, Ulrike Jahn, Eva Eylert, Ivan A. Berg, Daniel Kockelkorn, Wolfgang Eisenreich, and Georg Fuchs. A dicarboxylate/4-hydroxybutyrate autotrophic carbon assimilation cycle in the hyperthermophilic Archaeum Ignicoccus hospitalis. Proceedings of the National Academy of Sciences, 105(22):7851–7856, June 2008. Publisher: Proceedings of the National Academy of Sciences.

[10] Jan Zarzycki, Volker Brecht, Michael Müller, and Georg Fuchs. Identifying the missing steps of the autotrophic 3-hydroxypropionate CO_2_ fixation cycle in Chloroflexus aurantiacus. Proceedings of the National Academy of Sciences, 106(50):21317–21322, December 2009.

[11] Israel A. Figueroa, Tyler P. Barnum, Pranav Y. Somasekhar, Charlotte I. Carlström, Anna L. Engelbrektson, and John D. Coates. Metagenomics-guided analysis of microbial chemolithoautotrophic phosphite oxidation yields evidence of a seventh natural CO2 fixation pathway. Proceedings of the National Academy of Sciences, 115(1):E92–E101, January 2018. Publisher: Proceedings of the National Academy of Sciences.

[12] Tongxing Zhao, Yin Li, and Yanping Zhang. Biological carbon fixation: a thermodynamic perspective. Green Chemistry, 23(20):7852–7864, 2021.

[13] Sulamita Santos Correa, Junia Schultz, Kyle J. Lauersen, and Alexandre Soares Rosado. Natural carbon fixation and advances in synthetic engineering for redesigning and creating new fixation pathways. Journal of Advanced Research, 47:75–92, May 2023.

[14] Uri Barenholz, Dan Davidi, Ed Reznik, Yinon Bar-On, Niv Antonovsky, Elad Noor, and Ron Milo. Design principles of autocatalytic cycles constrain enzyme kinetics and force low substrate saturation at flux branch points. eLife, 6:e20667, February 2017. Publisher: eLife Sciences Publications, Ltd.

[15] P. Muller. Glossary of terms used in physical organic chemistry (IUPAC Recommendations 1994). Pure and Applied Chemistry, 66(5):1077–1184, January 1994. Publisher: De Gruyter.

[16] Jakob L Andersen, Christoph Flamm, Daniel Merkle, and Peter F Stadler. Defining Autocatalysis in Chemical Reaction Networks. Journal of Systems Chemistry, 8:121– 133, 2020.

[17] Shanshan Luo, Paul P. Lin, Liang-Yu Nieh, Guan-Bo Liao, Po-Wen Tang, Chi Chen, and James C. Liao. A cell-free self-replenishing CO2-fixing system. Nature Catalysis, 5(2):154–162, February 2022. Number: 2 Publisher: Nature Publishing Group.

[18] Thomas Schwander, Lennart Schada von Borzyskowski, Simon Burgener, Niña Socorro Cortina, and Tobias J. Erb. A synthetic pathway for the fixation of carbon dioxide in vitro. Science, 354(6314):900–904, November 2016. Publisher: American Association for the Advancement of Science.

[19] Richard McLean, Thomas Schwander, Christoph Diehl, Niña Socorro Cortina, Nicole Paczia, Jan Zarzycki, and Tobias J. Erb. Exploring alternative pathways for the in vitro establishment of the HOPAC cycle for synthetic CO2 fixation. Science Advances, 9(24):eadh4299, June 2023. Publisher: American Association for the Advancement of Science.

[20] Hannes Löwe and Andreas Kremling. In-Depth Computational Analysis of Natural and Artificial Carbon Fixation Pathways. BioDesign Research, 2021:9898316, September 2021. Publisher: American Association for the Advancement of Science.

[21] Jakob L. Andersen, Christoph Flamm, Daniel Merkle, and Peter F. Stadler. A Software Package for Chemically Inspired Graph Transformation. In Rachid Echahed and Mark Minas, editors, Graph Transformation, pages 73–88, Cham, 2016. Springer International Publishing.

[22] Jakob L. Andersen, Christoph Flamm, Daniel Merkle, and Peter F. Stadler. Chemical Transformation Motifs—Modelling Pathways as Integer Hyperflows. IEEE/ACM Transactions on Computational Biology and Bioinformatics, 16(2):510–523, March 2019. Conference Name: IEEE/ACM Transactions on Computational Biology and Bioinformatics.

[23] Arren Bar-Even, Elad Noor, Nathan E. Lewis, and Ron Milo. Design and analysis of synthetic carbon fixation pathways. Proceedings of the National Academy of Sciences, 107(19):8889–8894, May 2010.

[24] Minoru Kanehisa, Michihiro Araki, Susumu Goto, Masahiro Hattori, Mika Hirakawa, Masumi Itoh, Toshiaki Katayama, Shuichi Kawashima, Shujiro Okuda, Toshiaki Tokimatsu, and Yoshihiro Yamanishi. KEGG for linking genomes to life and the environment. Nucleic Acids Research, 36(Suppl_1):D480–D484, January 2008.

[25] Flamholz, E. Noor, A. Bar-Even, and R. Milo. eQuilibrator–the biochemical thermodynamics calculator. Nucleic Acids Research, 40(D1):D770–D775, January 2012.

[26] Moritz E Beber, Mattia G Gollub, Dana Mozaffari, Kevin M Shebek, Avi I Flamholz, Ron Milo, and Elad Noor. eQuilibrator 3.0: a database solution for thermodynamic constant estimation. Nucleic Acids Research, 50(D1):D603–D609, January 2022.

[27] Lu Xiao, Guoxia Liu, Fuyu Gong, Huawei Zhu, Yanping Zhang, Zhen Cai, and Yin Li. A Minimized Synthetic Carbon Fixation Cycle. ACS Catalysis, 12(1):799–808, January 2022.

[28] Claire R. Shen, Ethan I. Lan, Yasumasa Dekishima, Antonino Baez, Kwang Myung Cho, and James C. Liao. Driving Forces Enable High-Titer Anaerobic 1-Butanol Synthesis in Escherichia coli. Applied and Environmental Microbiology, 77(9):2905– 2915, May 2011. Publisher: American Society for Microbiology.

[29] Stefan Müller, Christoph Flamm, and Peter F. Stadler. What makes a reaction network “chemical”? Journal of Cheminformatics, 14(1):63, September 2022.

[30] Jakob L. Andersen, Christoph Flamm, Daniel Merkle, and Peter F. Stadler. Generic Strategies for Chemical Space Exploration, April 2014. 1302.4006 [cs, q-bio].

[31] Jakob L. Andersen and Daniel Merkle. A Generic Framework for Engineering Graph Canonization Algorithms. In 2018 Proceedings of the Meeting on Algorithm Engineering and Experiments (ALENEX), Proceedings, pages 139–153. Society for Industrial and Applied Mathematics, January 2018.

[32] Jeffrey D. Orth, Ines Thiele, and Bernhard Ø Palsson. What is flux balance analysis? Nature Biotechnology, 28(3):245–248, March 2010. Publisher: Nature Publishing Group.

[33] Gurobi Optimization, LLC. Gurobi Optimizer Reference Manual, 2024.

[34] Elad Noor, Hulda S. Haraldsdóttir, Ron Milo, and Ronan M. T. Fleming. Consistent Estimation of Gibbs Energy Using Component Contributions. PLOS Computational Biology, 9(7):e1003098, July 2013. Publisher: Public Library of Science.

[35] Matthew D. Jankowski, Christopher S. Henry, Linda J. Broadbelt, and Vassily Hatzimanikatis. Group Contribution Method for Thermodynamic Analysis of Complex Metabolic Networks. Biophysical Journal, 95(3):1487–1499, August 2008.

